# The Mla pathway promotes *Vibrio cholerae* re-expansion from stationary phase

**DOI:** 10.1101/2024.11.07.622497

**Authors:** Deborah R. Leitner, Franz G. Zingl, Alexander A. Morano, Hailong Zhang, Matthew K. Waldor

## Abstract

Bacteria have evolved diverse strategies to ensure survival under nutrient-limited conditions, where rapid energy generation is not achievable. Here, we performed a transposon insertion site sequencing loss-of-function screen to identify *Vibrio cholerae* genes that promote the pathogen’s fitness in stationary phase. We discovered that the Mla (<u>m</u>aintenance of lipid <u>a</u>symmetry) pathway, which is crucial for transferring phospholipids from the outer to the inner membrane, is critical for stationary phase fitness. Competition experiments with barcoded and fluorophore labeled wild-type and *mlaE* mutant *V. cholerae* revealed that the Mla pathway promotes re-expansion from 48h stationary phase cultures. The mutant’s defect in transitioning out of stationary phase into active growth (culturability) was also observed in monocultures at 48h. However, by 96h the culturability of the mutant and wild-type strains were equivalent. By monitoring the abundances of genomically barcoded libraries of wild-type and Δ*mlaE* strains, we observed that a few barcodes dominated the mutant culture at 96h, suggesting that the similarity of the population sizes at this time was caused by expansion of a subpopulation containing a mutation that suppressed the *mlaE* mutant’s defect. Whole genome sequencing revealed that *mlaE* suppressors inactivated flagellar biosynthesis. Additional mechanistic studies support the idea that the Mla pathway is critical for the maintenance of *V. cholerae’s* culturability as it promotes energy homeostasis, likely due to its role in regulating outer membrane vesicle shedding. Together our findings provide insights into the cellular processes that control re-expansion from stationary phase and demonstrate a previously undiscovered role for the Mla pathway.

**Importance:** Bacteria regularly encounter conditions with nutrient scarcity, where cell growth and division are minimal. Knowledge of the pathways that enable re-growth following nutrient restriction are limited. Here, using the cholera pathogen, we uncovered a role for the Mla pathway, a system that enables phospholipid re-cycling, in promoting *Vibrio cholerae* re-expansion from stationary phase cultures. Cells labeled with DNA barcodes or fluorophores were useful to demonstrate that though the abundances of wild-type and Mla mutant cells were similar in stationary phase cultures, they had marked differences in their capacities to regrow on plates. Of note, Mla mutant cells lose cell envelope components including high energy phospholipids due to OMV shedding. Our findings suggest that the defects in cellular energy homeostasis which emerge in the absence of the Mla pathway underlie its importance in maintaining *V. cholerae* culturability.

## Introduction

*Vibrio cholerae*, the causative agent of the diarrheal disease cholera, is a comma-shaped Gram-negative bacterium (1). Humans become infected via ingestion of contaminated food or water. Upon ingestion, *V. cholerae* colonizes the small intestine and expresses cholera toxin, often resulting in profuse secretory diarrhea, the hallmark of the disease. The diarrheal fluid releases the pathogen into the environment, where *V. cholerae* can reside for long periods in aquatic habitats and faces fluctuating challenges, including rapid changes in osmolarity and nutrient limitation (2, 3).

When nutrients are plentiful, bacteria can grow exponentially, and when nutrients become depleted, population growth arrest or “stationary phase” occurs (4, 5). As *V. cholerae* organisms enter stationary phase in lysogeny broth (LB), a typical rich medium, they undergo morphological and physiological changes in response to changes in their environment, particularly when the pH of the medium increases as bacteria shift from utilizing carbohydrates to amino acids for nutrition (6, 7). Prolonged cultivation in stationary phase is also known to select for ‘GASP’ (growth advantage in stationary phase) mutants that can often more efficiently scavenge residual nutrients (8–11). Interestingly, prolonged cultivation in stationary phase can lead *V. cholerae* to enter a state in which they are metabolically active but lose their ability to grow on routine culture media, the so-called ‘viable but non-culturable (VBNC)’ state (12–15).

The cell envelope of Gram-negative bacteria consists of two membranes - an inner membrane and an outer membrane - separated by the periplasm. While the inner membrane contains only phospholipids in both its leaflets, the outer membrane is comprised of phospholipids in its inner leaflet and predominately lipopolysaccharide (LPS) in its outer leaflet (16). During the transition from exponential to stationary phase, the composition of the cell envelope undergoes several changes that are thought to enhance stress resistance. For example, when *E. coli* enters stationary phase there is an increase in the thickness and cross-linking of the peptidoglycan layer, an overall decrease in the amount of protein in the outer membrane, and an increase in LPS in the outer membrane that changes the charge (17). However, knowledge of the processes that maintain cell envelope homeostasis in stationary phase have received relatively little attention.

Here, we carried out a transposon insertion site sequencing (Tn-seq) screen to identify *V. cholerae* genes that promote fitness in stationary phase LB cultures. Unexpectedly, Mla genes, which constitute an ABC transport system critical for <u>m</u>aintenance of lipid <u>a</u>symmetry in the Gram-negative cell envelope (18), were among the tops hits, revealing an unexplored link between cell envelope homeostasis and stationary phase fitness. Barcode- and fluorophore-labeled cells were used to compare the abundances of wild-type and *mlaE* mutant *V. cholerae* in stationary phase co-culture and monoculture. These studies revealed that the Mla pathway markedly altered the dynamics of *V. cholerae’s* culturability, specifically its capacity to return to growth from stationary phase.

## Results

### The Mla pathway promotes *V. cholerae* stationary phase fitness

To define the genes that could contribute to *V. cholerae* fitness under nutrient-limited conditions, we used Tn-seq to identify genes that promote *V. cholerae* fitness during stationary phase in LB culture. A transposon library created in a 2010 *V. cholerae* clinical isolate from Haiti (19) composed of ∼195,000 unique mutants was grown in a shaking LB culture at 37°C and samples were plated at 0, 8, 24 and 48h. Comparisons of the profiles between 8 and 24h (‘early stationary’) were used to identify genes that are important during early stationary phase (Table S1). Lastly, a comparison of the insertion landscapes between 24 and 48h (‘late stationary’) was used to gauge differences in the genetic requirements for viability and growth between early and late stationary phase (Table S1).

There were substantial differences in the number of genes that augment fitness (‘beneficial genes’, i.e. genes with fewer insertions, located on the left side of the volcano plot) between early and late stationary phase samples (Fig. 1AB, Table S1). In early stationary phase, 39 genes were found to promote fitness (4-fold change in abundance and Log_10_(1/p) value > 2) (Fig. 1A), while 285 genes met these criteria in late stationary phase (Fig. 1B). In contrast, the number of genes classified as ‘detrimental’ to fitness, with elevated abundance of insertions, was similar at 24 and 48h. Insertions in *rpoS*, were more abundant in late stationary phase samples (Fig. 1B, Table S1), as anticipated based on studies in *E. coli* (6, 20, 21), where reduced RpoS activity is a commonly observed adaptation in LB cultures (6).

**Fig. 1:**
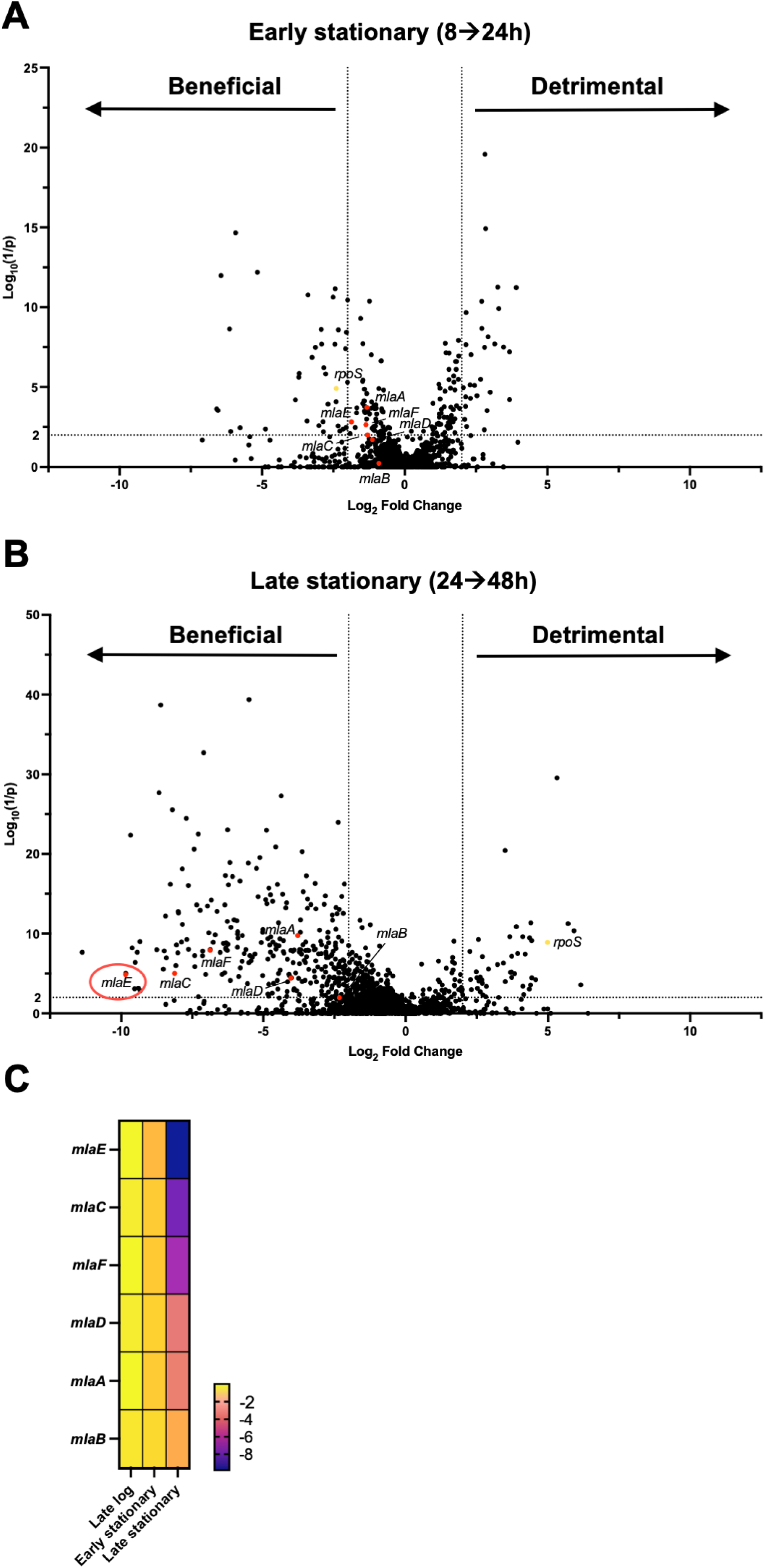
Transposon insertion sequencing identifies *V. cholerae* genes contributing to early and late stationary phase fitness. A-B) Transposon screen of early (A) and late (B) stationary phase WT cultures. The X-axis shows Log_2_ fold change in insertion frequency, the Y-axis shows corresponding inverse *P*-values from the Mann–Whitney U test. Points represent genes. Genes shown as red dots are those involved in the Mla pathway; *rpoS* is shown as a yellow dot. C) Differences in the insertion frequency in genes involved in the Mla pathway from WT cultures grown to late log phase (0-8h), early (8-24h) and late stationary phase (24-48h); a darker color indicates a greater negative fold change and thus fewer insertions.

Among the most prominent pathways whose disruption decreased fitness in comparisons between the 24 and 48h samples were in genes of the Mla pathway, which is involved in maintaining the lipid asymmetry of the outer membrane. This highly conserved pathway in Gram-negative bacteria consists of an outer membrane lipoprotein, MlaA, a periplasmic lipid carrier, MlaC, and the inner membrane transporter complex, MlaBDEF, that enables retrograde transport of phospholipids from the outer to the inner membrane (18). Deletion of any of the Mla genes disrupts this system, leading to an accumulation of phospholipids in the outer leaflet of the outer membrane and elevated loss of outer membrane components (phospholipid, LPS, and proteins) via outer membrane vesicles (OMVs) (22). In the case of *V. cholerae*, this enables rapid alteration of the bacterial membrane composition resulting in faster adaption to the host environment (23).

In early stationary phase *V. cholerae* cultures, insertions in Mla genes were not significantly decreased compared to late exponential phase (Fig. 1A), but at 48h disruption of *mlaACDEF* had decreased significantly, indicating that Mla genes are beneficial in late stationary phase (Fig. 1BC). At this point, the fitness defect of *mlaE* insertions were particularly striking, with a ∼1,000-fold reduction in insertions between 24 and 48h (Fig. 1BC).

### *mlaE* modifies *V. cholerae* culturability in stationary phase

A Δ*mlaE* mutant was created to validate the Tn-seq results and to determine whether *mlaE* deficiency reduces *V. cholerae* viability and/or culturability in 48h stationary phase cultures. The mutant and WT *V. cholerae* were competed in co-culture experiments (Fig. 2A). Three unique ∼20 nucleotide barcodes were integrated into the WT and the Δ*mlaE* mutant at a neutral genomic locus, enabling differentiation and quantification of the abundance of the two strains in co-culture by barcode sequencing (24). The barcoded mutant and WT strains were separately grown overnight and then mixed in a 1:1 ratio. Every 24h, aliquots were removed from the cultures and the barcodes were either directly sequenced (direct) or plated and grown (outgrowth) before barcode sequencing (Fig. 2A). The STAMPR analysis pipeline (24, 25) was used to determine the relative abundance of each barcoded strain within the co-culture, enabling calculation of a competitive index (CI). When samples were taken directly (no outgrowth on plates) from the co-culture, there was no difference in the abundance between WT and Δ*mlaE* strains after 24h, but the mutant had a slight fitness defect after 48h (Fig. 2B). In contrast, when samples were plated prior to sequencing (outgrowth), at the 24h timepoint, the Δ*mlaE* strain had a ∼3-fold fitness defect. At 48h, the fitness disadvantage of the Δ*mlaE* mutant after plating was much more pronounced (∼830-fold) (Fig. 2B). These observations align with the 48h Tn-seq findings, where there was a striking reduction in the abundance of insertions in *mlaE*.

**Fig. 2:**
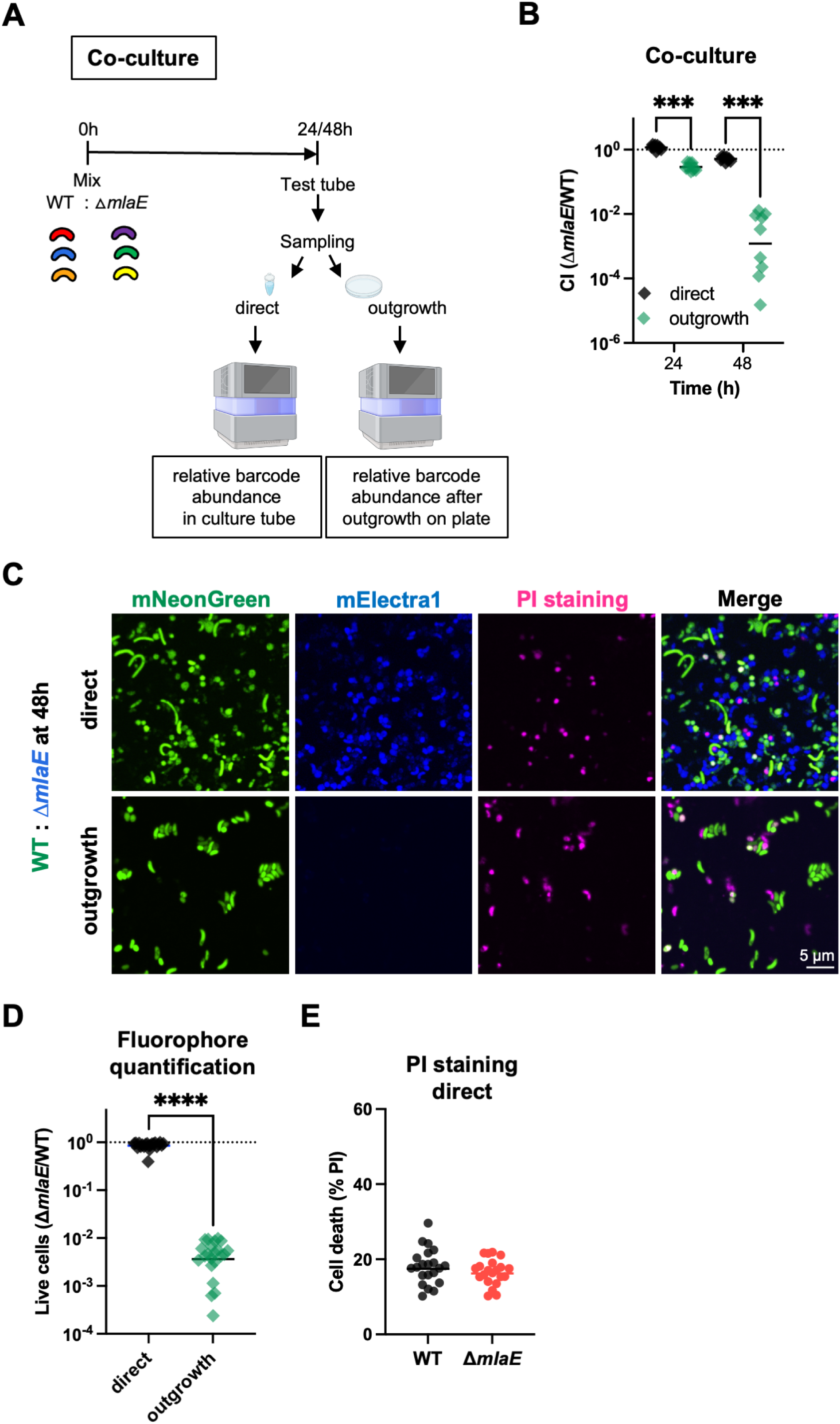
*mlaE* modifies *V. cholerae* culturability in stationary phase. A) Diagram of experimental scheme. Barcoded WT and Δ*mlaE* strains, each containing 3 unique barcodes, were used in co-culture experiments. After 24 and 48h, aliquots were either directly sequenced (direct) or plated on LB plates (outgrowth) before sequencing. B) Competitive indices (*ΔmlaE*/WT) of co-culture experiments using barcoded WT and mutant strains. The relative abundance of each strain was measured by amplicon sequencing. Graph depicts results of 3 independent experiments; ***P < 0.001 Kruskal-Wallis test, Multiple comparisons. C) Representative images of co-cultures of mNeonGreen-labeled WT (green) and mElectra1-labelled Δ*mlaE* (blue); PI (pink) was used to detect dead cells. D) Quantification of fluorophore-positive WT and mutant cells at 48h. (n=20 fields; ****P < 0.0001; Mann-Whitney test). E) Quantification of dead (PI-positive) WT and Δ*mlaE* cells at 48h. n=20 fields. For all graphs, lines represent geometric means and significant differences between data sets are marked by asterisks.

Fluorescence microscopy was used as alternative approach to quantify the abundance of the co-cultured WT and Δ*mlaE* strains (Fig. 2C). The ratios of green fluorophore-labelled WT cells to blue fluorophore-labelled Δ*mlaE* cells were similar when samples were taken directly from a 48h culture (Fig. 2CD). There was no difference in the fraction of dead cells (measured as cells staining with propidium iodide (PI)) between the two strains (Fig. 2E). Interestingly, there was a modest difference in the shape of the two strains; WT cells had a higher number of elongated cells compared to Δ*mlaE* cells (Fig.2C). Very similar findings were obtained when the fluorophore labelling was switched (WT: blue and Δ*mlaE*: green) (Fig. S1) indicating that the labelling did not impact the conclusions. In marked contrast, when samples from the 48h culture were plated and grown until colonies formed, there were at least 300-fold greater WT than Δ*mlaE* cells (Fig. 2D). These observations mirror the barcode-based quantification and support the idea that the Δ*mlaE* strain has a diminished capacity to re-grow after reaching stationary phase at 48h.

We used stationary phase grown monocultures of the WT and Δ*mlaE* strains to further investigate the culturability defect of the mutant. The two strains were grown separately in LB at 37°C with shaking for 96h. Every 24h, aliquots were removed to measure optical densities and colony-forming units (CFU). The OD_600_ measurements of both cultures were indistinguishable throughout the experiment but the CFU dynamics differed (Fig. 3A). At 24h, the WT and Δ*mlaE* strains had similar CFU, whereas at 48h, the WT had ∼28-fold more (p=0.000155) CFU compared to the mutant (Fig. 3A). Both the similarity of the OD_600_ measurements and the 28-fold difference in the CFU of the two cultures at this point strongly suggests that the Δ*mlaE* mutant has a defect in its culturability on solid media. The magnitude of the defect was more pronounced in co-culture experiments, where there was ∼1,000-fold lower abundance in Δ*mlaE* derived barcodes compared to those from WT at 48h (Fig. 2B), suggesting that interactions between the two strains in co-culture may further impair the mutant’s viability and/or culturability.

**Fig. 3:**
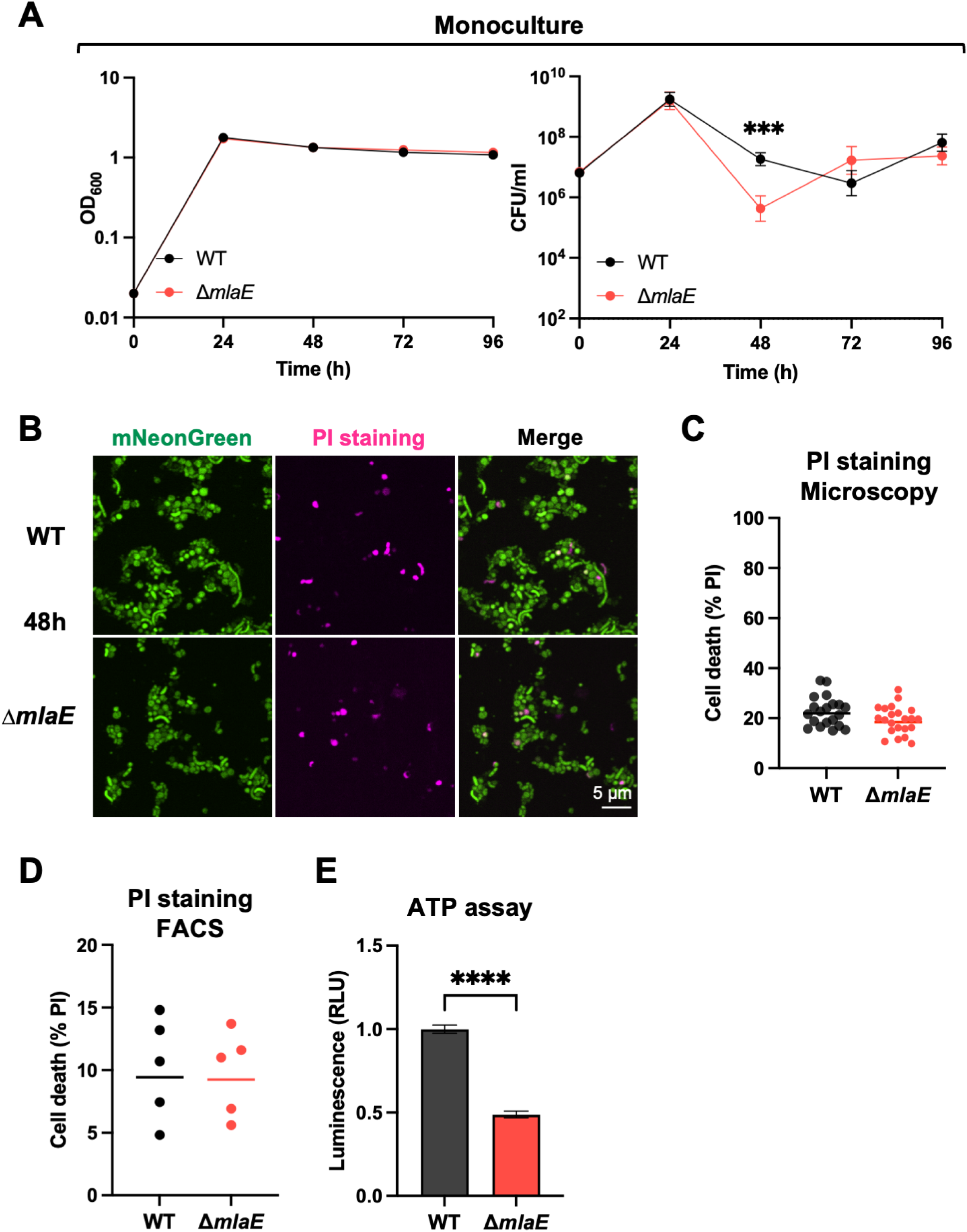
Δ*mlaE V. cholerae* are viable but less metabolically active after 48h. A) Optical densities and CFU/ml of WT and *ΔmlaE* monocultures over 96h of continuous culture. The lines representing OD_600_ measurements shown on the right are geometric means. Graph depicts results of 8 independent experiments; P = 0.000155; Multiple Mann-Whitney tests. B) Representative images of 48h monocultures of mNeonGreen-labeled WT and mNeonGreen-labeled Δ*mlaE;* PI (pink) was used to detect dead cells. C) Quantification of dead (PI-positive) WT and Δ*mlaE* cells after 48h (n=20 fields). D) Quantification of dead (PI-positive) WT and Δ*mlaE* cells by FACS analysis. Graph depicts results of 5 independent experiments. E) Cellular ATP levels of WT and Δ*mlaE* monocultures after 48h. Graph depicts results of 10 independent experiments; ****P < 0.0001; Mann-Whitney test. For all graphs, lines represent geometric means, error bars represent standard deviations and significant differences between data sets are marked by asterisks.

Imaging and FACS analysis were also used to characterize the viability of Δ*mlaE* cells taken directly from 48h monocultures. The fraction of dead cells, determined by PI staining was similar in both monocultures at 48h (Fig. 3C). FACS analysis corroborated the latter result, as a similar percentage of PI- positive dead cells was observed in five independent cultures of both strains (Fig. 3C). The relative cellular ATP levels in the two 48h monocultures was also assessed as a measure of metabolic activity in both strains. Notably, WT cells had ∼2x higher ATP levels compared to mutant cells (Fig. 3E), suggesting that reduced energy availability might contribute to the observed loss of culturability in the *mlaE* mutant.

### Characterization of Δ*mlaE* suppressor mutations

Interestingly, the Δ*mlaE* mutant’s impaired culturability relative to the WT did not persist. The CFU in the Δ*mlaE* culture increased from 48 to 72h post inoculation, whereas the WT CFU decreased during this interval. At 96h, the CFU in both cultures were similar (Fig. 3A). We hypothesized that the similar CFU of the WT and Δ*mlaE* strains at 96h, despite the 28-fold fewer Δ*mlaE* CFU at 48h (Fig. 3A), represents the emergence of suppressor mutants in the Δ*mlaE* culture. Barcoded libraries, each including >50,000 unique barcodes of the WT and Δ*mlaE* strains were used to monitor the dynamics of subpopulations within monocultures (Fig. 4A). Every 24h, aliquots from the cultures were plated and the barcode abundances were determined by sequencing (Fig. 4A). We reasoned that barcodes in bacteria with Δ*mlaE* suppressor mutations that promote fitness in stationary phase should become more abundant in the culture, particularly after 48h, when the CFU of the Δ*mlaE* culture increased (Fig. 3A).

**Fig. 4:**
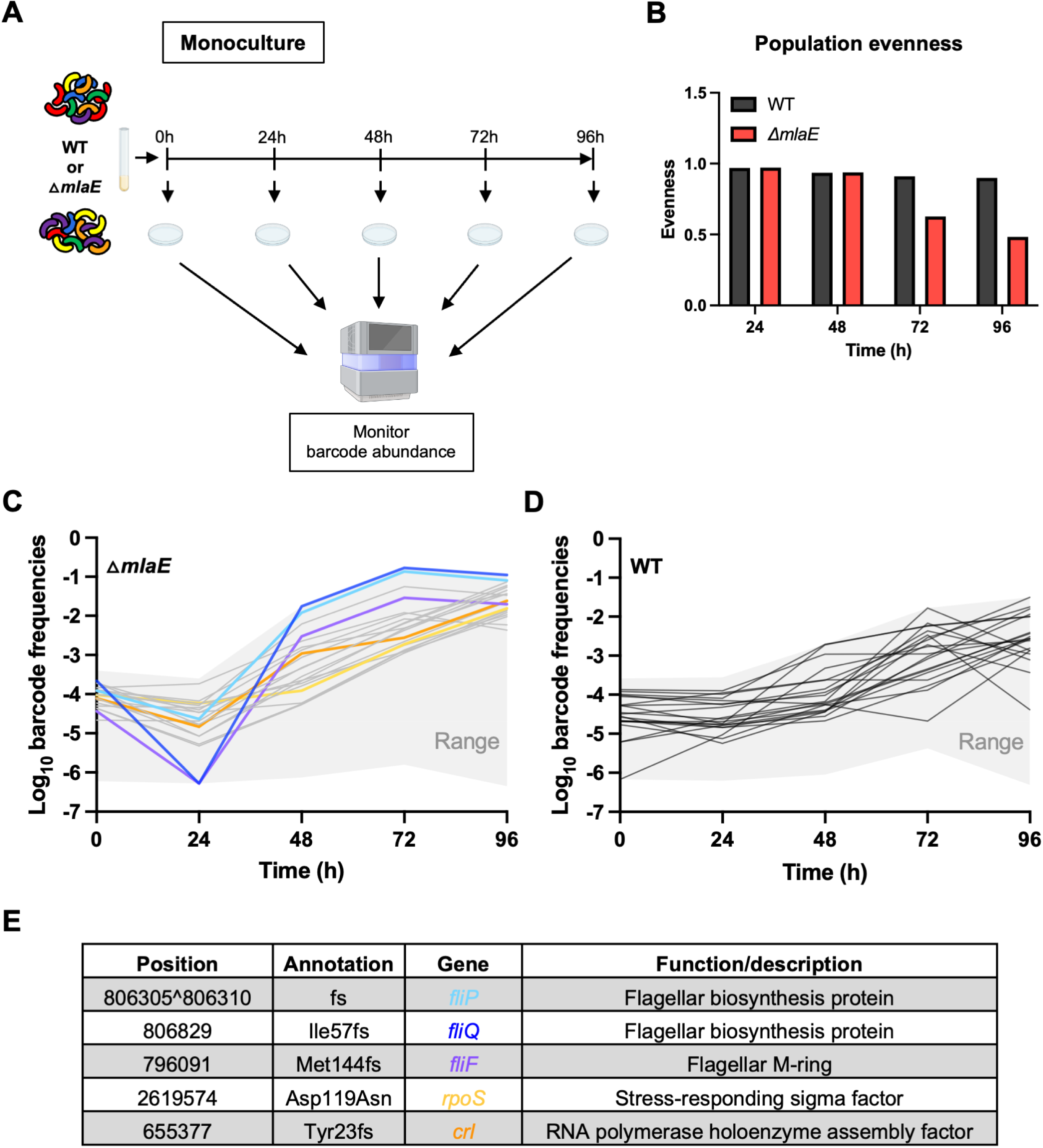
Detecting and tracing suppressor lineages in Δ*mlaE* cultures with barcodes. A) Diagram of experimental scheme. Monocultures of WT and Δ*mlaE* barcoded libraries were grown for 96h. Every 24h, aliquots were plated for sequencing to determine the relative abundances of the barcodes in the culture. B) Population evenness of the barcodes (lineages) of WT and mutant strains over time, calculated using the Shannon index. C-D) Population dynamics of barcoded WT and Δ*mlaE* strains were studied over 96h. The relative abundance of each strain was measured by amplicon sequencing. Trajectories of the top 20 barcodes of the mutant (C) and WT (D) are shown as colored and grey lines respectively. The grey area depicts the range of all barcode trajectories of the respective strain. Highlighted in C are the lineage trajectories of Δ*mlaE fliP\** in light blue; Δ*mlaE fliQ\** in dark blue; Δ*mlaE fliF\** in purple; Δ*mlaE rpoS\** in yellow and Δ*mlaE crl\** in orange. The asterisks (*) represent the spontaneous suppressor mutations. E) Mutations present in the characterized suppressor mutants appearing in Δ*mlaE* after 96h along with a description of the function of each gene. fs, frameshift.

The trajectories of barcode frequencies in the WT and the Δ*mlaE* cultures differed. The distribution of the barcode abundances (population evenness) in the two cultures was similar at 24 and 48h, but not at 72h, suggesting that several barcodes in the Δ*mlaE* population increased in their frequency (Fig. 4B). At 48h, the 20 most abundant barcodes in the Δ*mlaE* culture represented 5.4% of the culture, whereas the 20 most abundant barcodes in the WT culture represented only 1.1% of the culture at this point. By 96h, the most abundant barcodes in the Δ*mlaE* culture became even more dominant; the two most abundant barcodes represented 19.3 % of the culture and the top 20, comprised 69.6% (Fig. 4C). In the WT culture at this point, the 20 most abundant barcodes represented only 12.8% of the population (Fig. 4D).

To identify potential suppressor mutations that would likely account for the elevated abundance of a subset of barcodes, colonies from the plated 96h Δ*mlaE* culture were randomly picked for sequencing of their barcodes; if their barcodes were among the abundant barcodes, their genomes were sequenced. The two most abundant barcodes, which together represented ∼19% of the Δ*mlaE* population at 96h (Fig. 4C, blue barcodes), contained mutations in *fliP* and *fliQ* (Fig. 4E). The products of these two genes are both membrane proteins that form a complex with FlhA, FlhB and FliR in the central pore (‘export gate’) of the MS ring (26) (Fig. S2). FliF, which anchors the flagellum in the cytoplasmic membrane, (purple in Fig. 4C, Fig. S2) was also among the abundant barcodes. Thus, inactivation of flagellar biosynthesis appears to represent a means to bypass the culturability defect of the Δ*mlaE* strain. Along with these flagellar biosynthesis genes, we identified putative suppressor mutations in *rpoS* (yellow barcode), and *crl* (orange barcode) (Fig. 4CE), a gene implicated in RpoS activity (27). Inactivation of both *rpoS* and *crl* have been shown to promote stationary phase fitness (6, 28). The orange and yellow barcodes, which track the lineages with mutations in *rpoS* and *crl*, gradually became more abundant from ∼48- 96h, whereas the most prominent change in the abundance of the three flagellar mutants occurred from 24-48h.

### Defective cellular energy homeostasis may account for the culturability defect of Δ*mlaE V. cholerae*

We used the spontaneously arising Δ*mlaE fliP*,* Δ*mlaE fliQ\** and Δ*mlaE fliF\** mutant strains to investigate how mutations in these flagellar biosynthesis genes suppress the culturability defect of the Δ*mlaE* strain. Initially, we corroborated that after 48h monocultures, all three of these strains had elevated CFU compared to the Δ*mlaE* mutant (Fig. 5A), suggesting that these mutations elevate the stationary phase fitness of the Δ*mlaE* mutant. All three of these strains had severe motility defects in soft agar and lacked a flagellum (Fig. 5BC). Since we found that the Δ*mlaE* strain had reduced ATP content in 48h monocultures (Fig. 3E), we hypothesized that the absence of the energy expenditure required for flagellar biosynthesis and/or spinning, could result in elevations in cellular ATP content in these strains. This prediction proved correct, as all three strains had ∼50% higher ATP content than the Δ*mlaE* strain but less than the WT (Fig. 5D). Since the Δ*mlaE* strain had elevated OMV shedding and concomitant phospholipid loss, another potential mechanism that the flagellar mutants could have restored the culturability of the Δ*mlaE* mutant was if the absence of the flagellum reduced OMV shedding; however, all suppressor mutants released similar OMV amounts compared to the Δ*mlaE* strain they were derived from (Fig. S3). Together, these observations suggest that the conservation of cellular ATP content by the Mla pathway may be a critical mechanism through which this phospholipid recycling pathway maintains the culturability of *V. cholerae* in stationary phase. To further test this idea, we grew several WT and Δ*mlaE* cultures for 48h, supplementing half of the cultures with 0.2% glucose after 24h (Fig. 5E). The addition of glucose as an energy source yielded higher CFU counts in both strains compared to the LB only control. Notably, glucose supplementation eliminated the culturability defect of the Δ*mlaE* mutant, and the CFU in the WT and Δ*mlaE* cultures were similar (Fig. 5F). Furthermore, glucose supplementation elevated the cellular ATP content in the mutant, from 50% of WT to ∼82% of WT with glucose (Fig. 5G). Collectively, these findings support the idea that the culturability defect of the Δ*mlaE* strain results from depletion of cellular energy levels.

**Fig. 5:**
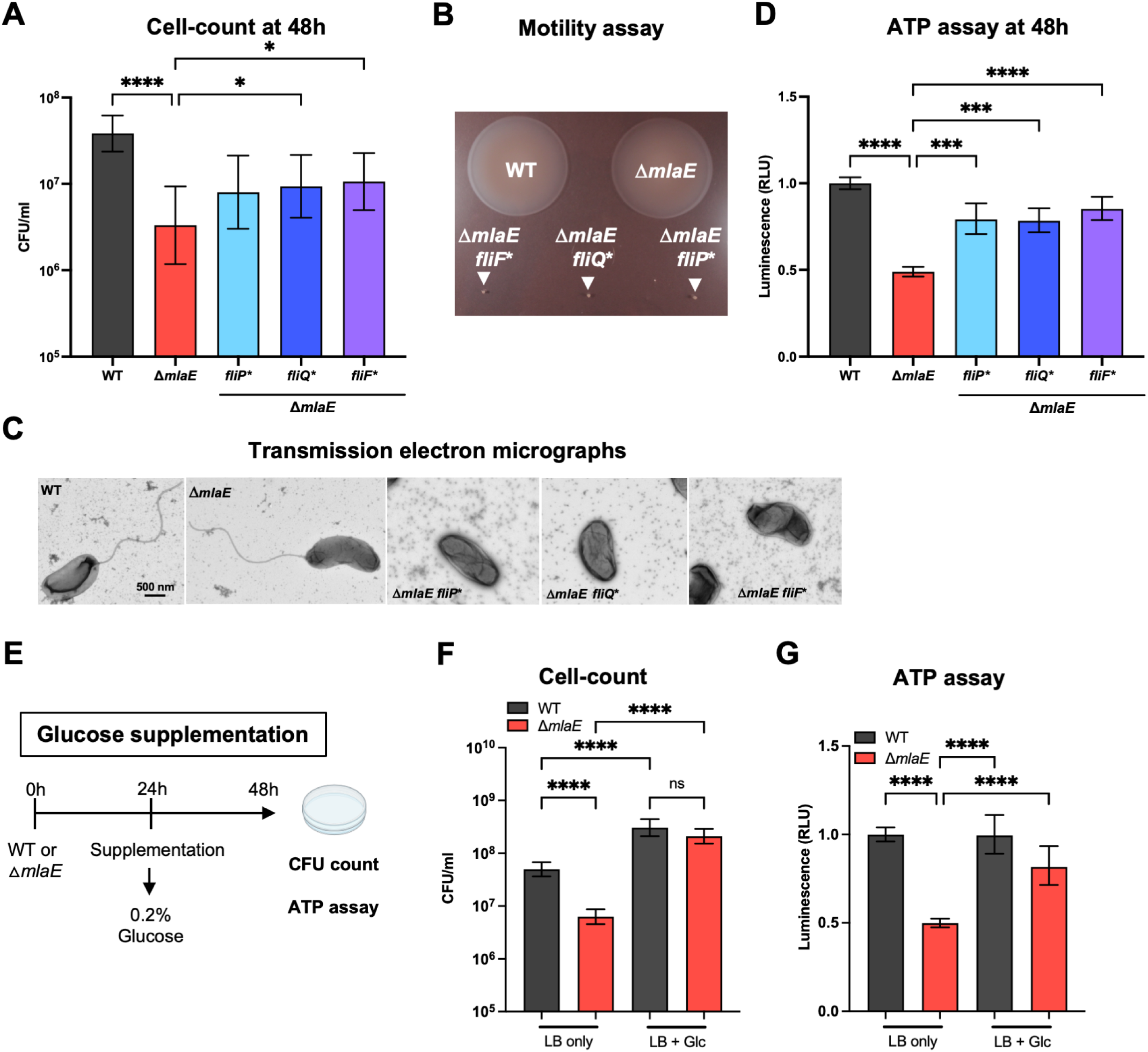
Flagellar mutants partially restore cellular ATP levels and rescue the culturability defect of Δ*mlaE V. cholerae*. A) CFU counts of WT, Δ*mlaE,* Δ*mlaE fliP*,* Δ*mlaE fliQ* and* Δ*mlaE fliF\** strains after 48h in LB culture. Graph depicts results of ≥8 experiments; *P < 0.05; ****P < 0.0001 (Kruskal-Wallis test). B) Swarming motility of WT, Δ*mlaE,* Δ*mlaE fliF*,* Δ*mlaE fliQ* and* Δ*mlaE fliP\** strains after 20h of incubation in soft agar. C) Transmission electron micrographs of WT, Δ*mlaE,* Δ*mlaE fliP*,* Δ*mlaE fliQ* and* Δ*mlaE fliF\** strains grown overnight in LB. D) Cellular ATP levels of WT, Δ*mlaE,* Δ*mlaE fliP*,* Δ*mlaE fliQ* and* Δ*mlaE fliF\** cultures after 48h. Graph depicts results of ≥6 independent experiments; ***P < 0.001; ****P < 0.0001 (Brown-Forsythe and Welch ANOVA tests). The data sets of the WT and Δ*mlaE* are also presented in figure 3E. E) Diagram of experimental scheme. Monocultures of WT and Δ*mlaE* strains were grown for 48h. After 24h, half of the cultures were supplemented with 0.2% glucose. After a total of 48h, supplemented and non- supplemented cultures of both strains were plated for CFU counts and cellular ATP levels were measured. F) CFU counts of WT and Δ*mlaE* with and without glucose supplementation. Graph depicts results of 10 independent experiments; ****P < 0.0001 (Brown-Forsythe and Welch ANOVA tests). G) Cellular ATP measurement of WT and Δ*mlaE* cultures with and without glucose supplementation. Graph depicts results of ≥8 independent experiments; ****P < 0.0001 (Brown-Forsythe and Welch ANOVA tests). For all graphs, lines represent geometric means, error bars represent standard deviations, and significant differences between data sets are marked by asterisks.

## Discussion

Here, a Tn-seq screen uncovered an uncharacterized link between the Mla pathway and *V. cholerae* stationary phase fitness. Co-culture and monoculture experiments using fluorophore-labeled or barcoded versions of the WT and Δ*mlaE* strains revealed that the Mla pathway had a marked influence on the capacity of this pathogen to resume growth (culturability) from stationary phase but only a modest influence on its apparent persistence in liquid culture. These findings highlight the utility of DNA barcodes and fluorophores for cell culture independent quantification of bacterial abundance. Although WT and Δ*mlaE* cultures had similar OD_600_ measurements after 48h, there were 28-fold fewer Δ*mlaE* vs WT CFU recovered on plates (Fig. 3A). Both live-dead staining and fluorescence microscopy of fluorophore-bearing cells suggested that at 48h the Δ*mlaE* cells were in a ‘viable but non-culturable state’. After 96h of monoculture, the abundances of the WT and Δ*mlaE* strains were similar after plating, suggesting that the mutant regained its culturability. Creation of barcoded libraries of the WT and Δ*mlaE* strains enabled facile characterization of both strains’ population trajectories over time in culture. In the Δ*mlaE* culture, mutants in flagellar biosynthesis arose by 48h and became dominant in the population. Additional studies supported the idea that cellular ATP levels exert potent control over *V. cholerae* culturability. Since Mla mutants lose phospholipids and other high energy outer membrane components due to increased shedding of OMVs, we propose that the defects in cellular energy homeostasis that emerge in the absence of the Mla pathway underlie its role in maintaining *V. cholerae* culturability.

The application of high throughput DNA sequencing to quantify DNA barcodes in microbial populations represents a robust method for tracking the abundance of microbial cells, tracing lineages in various contexts, and for quantifying infection bottlenecks (25, 29–36). Our study underscores the value of using sequencing and barcodes to quantify cell abundance independent of cell culture methods thereby facilitating investigation of determinants of culturability. Furthermore, by tracking the dynamics of the barcoded Δ*mlaE* strain, we were able to distinguish between two possible hypotheses for why the mutant’s culturability was restored at 96h. In principle, either *a)* a large proportion of previously non-culturable cells could have become culturable again or *b)* a small fraction of suppressor mutants could have outgrown the existing population, increasing the mutant strain’s culturability. By analyzing barcode frequencies over time, we proved the second hypothesis correct and found that a few dominant barcodes (lineages), which harbored mutations inactivating flagellar biosynthesis, emerged in the Δ*mlaE* population at 48h, with the two most abundant barcodes representing ∼20 % of the population by 96h.

While extensive studies of the changes in bacterial physiology that accompany nutrient starvation and growth arrest in stationary phase have been carried out (4), there has been limited application of Tn-seq in this research area. Our Tn-seq screen revealed that insertions in *rpoS* and flagellar genes (Fig. 1B and Table S1) became more abundant after 48h, indicating that these genes impair stationary phase fitness in *V. cholerae*, as has been seen in previous studies of other organisms (6, 8, 37).

Notably, our screen revealed that insertions in the genes of the Mla pathway became depleted by 48h, suggesting that this pathway is critical for *V. cholerae* fitness in stationary phase. A recent proteomics-based study of *E. coli* survival in stationary phase also uncovered that many cell envelope localized proteins, including components of the Mla pathway, are particularly important for survival during long-term starvation (38).

The Mla pathway is a highly conserved phospholipid transport system in Gram-negative bacteria that maintains outer membrane asymmetry (18). Disruptions in this system lead to phospholipid accumulation in the outer membrane that is resolved by an increased release of OMVs (22). Inactivation of the Mla pathway in exponentially growing *V. cholerae* enables rapid alteration of the membrane composition due to OMV shedding, resulting in faster adaption to the host environment (23). In contrast to the beneficial effect of *mlaE* inactivation in log phase, deletion of *mlaE* significantly reduced cell culturability in late stationary phase. In *E. coli*, Mla null mutations also lead to hypervesiculation (22). However, unlike *V. cholerae*, *E. coli* has two additional systems, PldA and PagP, to ensure that phospholipids do not accumulate in the outer leaflet of the outer membrane (18). *V. cholerae* lacks homologs of both PldA and PagP and apparently relies solely on its Mla pathway for this function. Consequently, disruptions in the Mla system in *V. cholerae* lead to a distinct phenotype as phospholipids accumulate in the outer membrane and are shed via OMVs.

Our observations suggest that loss of energy-rich molecules like phospholipids, whose biosynthesis requires >20% of cellular energy (39), via OMVs in the absence of a functional Mla system are especially costly during nutrient limiting conditions.

After 48h of monoculture, the point at which a large proportion of Δ*mlaE* cells from liquid culture were no longer capable of re-growth on solid media, there was a marked reduction in cellular ATP in the *mlaE* mutant compared to the WT. Furthermore, the predominant suppressor mutations that arose in the Δ*mlaE* culture blocked biosynthesis of the flagellum and partially restored cellular ATP. Although flagellar motility often confers benefits, flagellar synthesis and assembly requires significant energy expenditure. In *E. coli,* it is estimated that the combined costs of constructing and operating flagella require ∼10% of the cell’s energy budget (40). Unlike *E. coli*, *V. cholerae’s* single polar flagellum is sheathed by the cell envelope, and it seems likely that substantial energy conservation is achieved by blocking flagellar synthesis. We speculate that the increase in cellular ATP levels in the absence of flagellum production and flagellar-based motility contribute to the increased culturability in the Δ*mlaE fliP\**, Δ*mlaE fliQ\** and the Δ*mlaE fliF\** mutants. Consistent with this hypothesis, a recent study reported that non-motile *V. cholerae* mutants remain culturable for longer periods compared to their motile parent strain (41). Further buttressing the idea that the culturability defect of the Δ*mlaE* mutant is linked to its lower cellular ATP levels, we found that provision of glucose to the Δ*mlaE* culture at 24h increased ATP levels and restored culturability on solid media. However, the mechanism(s) by which glucose supports culturability of stationary phase Δ*mlaE* cells requires further study. Nevertheless, it is tempting to speculate that cellular energy levels may underlie many cases of culturability defects observed with so-called VBNC phenotypes. Supporting this idea, a recent study in *E. coli* 0157:H7 found that higher cellular ATP levels correlated with faster resuscitation from the VBNC state in a mutant defective in LPS synthesis (Δ*rfaL*) (42).

In sum, our findings illustrate that labeling bacteria with DNA barcodes and fluorophores to monitor bacterial cells in stationary phase can be valuable for distinguishing whether stationary phase fitness defects result from cell death or deficiencies in re-growth in rich media. Deeper understanding of the processes and pathways that enable re-growth from nutrient deficient environments will offer insights into how pathogens that persist in the environment, such as *V. cholerae*, can give rise to infections.

## Materials and Methods

### Bacterial strains and growth conditions

Bacterial strains and oligonucleotides used in this study are listed in Table S2 and S3. All *V. cholerae* strains used in this study are derivatives of a spontaneous streptomycin (Sm)-resistant variant of a clinical isolate from the 2010 Haiti cholera outbreak (HaitiWT) (19). *E. coli* strain MFDλpir was used for genetic manipulations. All bacteria were grown in lysogeny broth (LB) broth or on LB agar plates with aeration at 37°C. If required, antibiotics, or other supplements were used in the following final concentrations: streptomycin (Sm), 200 μg/ml; carbenicillin (Cb), 100 μg/ml; kanamycin (Km), 50 μg/ml; diaminopimelic acid (DAP) 0.3 mM; glucose (Glc), 0.2%; sucrose (Suc), 10% and glycerol 35%.

### Strain and plasmid construction

Allelic exchange using derivatives of the suicide vector pCVD442 and the MFDλpir *E. coli* strain as donor, was used to construct in-frame *V. cholerae* deletion mutants as described (43); correct deletions were confirmed by PCR. *V. cholerae* codon-optimized versions of a blue (mElectra1) (44) or green (mNeonGreen) (45) fluorophore under control of the constitutive pTac promoter were created and introduced into pSM1 between the left and right repeats of a site specific Tn7 transposon (24). Triparental conjugations using either WT or Δ*mlaE V. cholerae* as recipients along MFDλpir *E. coli* containing the fluorophore-bearing plasmids or Tn7 transposase-containing ‘helper’ plasmid pJMP1039 (46) were used to create fluorescent derivatives of WT and Δ*mlaE V. cholerae*. These strains were verified by both PCR and fluorescence microscopy.

### Transposon insertion site sequencing

2 mL LB cultures were inoculated with the HaitiWT Himar transposon library (47) and grown at 37°C with shaking at 200 rpm. Samples were removed from the culture at 8, 24 and 48h and plated on 150 mm LB Sm plates. Cells were scraped up in LB glycerol and stored at -80°C. Genomic DNA was extracted using the GeneJet gDNA Isolation Kit and DNA libraries were prepared as described previously (48). Reads were analyzed using the R-based TnSeq analysis pipeline RTISAn (35). To identify over-or underrepresented loci, we used the 8h sample as the “input” and the 24 or 48h sample of the respective strain as the “output”.

### CFU plating and OD_600_ measurement

Overnight cultures of the WT and Δ*mlaE* strains were back diluted to an OD_600_: 0.01 and incubated at 37°C with shaking at 200 rpm until reaching the exponential phase (OD_600_: 0.2). The exponentially growing monocultures were diluted (1:10) and incubated at 37°C with shaking at 200 rpm for 96h. Serial dilutions of the respective strains were plated every 24h on LB Sm plates to determine viable counts. Additionally, the OD_600_ was measured in parallel every 24h using a BioTek Epoch2 microplate reader.

### Construction of barcoded *V. cholerae* libraries

Barcoded libraries of WT and **Δ***mlaE V. cholerae* were constructed as described (24). Briefly, WT and **Δ***mlaE V. cholerae* were conjugated with the *E. coli* donor strains MFDλpir pSM1 (barcode donor) and MFDλpir pJMP1039 (containing the helper Tn7 transposase). Transconjugants were selected on LB Sm Km plates, pooled in LB glycerol, and stored at -80°C in aliquots.

### Barcode sequencing and analysis

Barcode sequencing and analysis was essentially performed as previously described (24, 25). To amplify the barcoded region, samples were boiled and used for PCR. Amplicons were verified by gel electrophoresis, cleaned using the Qiagen PCR Purification kit, and sequenced on a NextSeq1000. Sequencing reads were processed using the STAMPR pipeline (demultiplexing, trimming, mapping and counting) to determine barcode frequencies (24, 25).

### Competition assays

Overnight cultures of the WT and **Δ***mlaE* (each containing 3 barcodes) *V. cholerae* were back diluted to an OD_600_: 0.01 and incubated at 37°C with shaking at 200 rpm until reaching the exponential phase (OD_600_: 0.2). The exponentially growing monocultures were mixed in a 1:1 ratio and used to inoculate cultures of 2 ml LB cultures. Co-cultures were incubated at 37°C with shaking at 200 rpm for 48h. Samples for barcode sequencing were either directly taken (2 µl of the co-culture in 100 µl ddH_2_O and boiled) after 24 and 48h or plated (outgrowth) on LB Sm plates. Bacteria were harvested form the LB Sm plates the next day and samples were processed and analyzed for barcode sequencing as described (24, 25). Competitive indices (CI) were calculated as the ratio of mutant to WT bacteria normalized to the input ratio. If no reads were detected, we substituted the value with 1, which corresponds to the limit of detection.

### Spontaneous suppressor isolation and sequencing

Overnight cultures of barcoded WT and **Δ***mlaE V. cholerae* were back diluted to an OD_600_: 0.01 and incubated at 37°C with shaking at 200 rpm until reaching the exponential phase (OD_600_: 0.2). The exponentially grown monocultures were diluted (1:10) and incubated at 37°C with shaking at 200 rpm for 96h. Serial dilutions of the respective cultures were plated every 24h on LB Sm plates to determine cell counts. Additionally, 50 µl of the respective samples were plated for barcode sequencing. To display the trajectories of the barcoded libraries, barcode frequencies of the respective samples were determined as previously described (24, 25). The population evenness was calculated using the Shannon index (49).

To isolate suppressor mutants in the Δ*mlaE* background, the 96h sample was re-streaked on a LB Sm plate. Single colonies were grown overnight, and their barcoded region was amplified. Strains that showed an increase in their barcode frequencies at 48h or 96h were subjected to whole genome sequencing (SeqCenter, Pittsburgh). Genome assembly and variant identification was performed using CLC Genomics Workbench 12 (Qiagen, Germany). Mutations were identified using the HaitiWT genome (50) as a reference and assigned as likely suppressors if they were present in >93% of reads.

### Microscopy

Samples from cultures of WT and Δ*mlaE V. cholerae* marked with either the mNeonGreen or the mElectra1 fluorophore (or vice versa) grown for 48h were immobilized on 0.8% agar pads containing a 1:1000 dilution of propidium iodide (PI) and imaged with a Nikon Ti2 Eclipse spinning disk confocal microscope using a 100x oil immersion lens with a numerical aperture of 1.45 and an Andor Zyla 4.2 Plus sCMOS monochrome camera. Image analysis was performed on ImageJ (FIJI (2.14.0) software using custom macros.

Samples from co-cultures of WT and Δ*mlaE V. cholerae* marked with either the mNeonGreen or the mElectra1 fluorophore (or vice versa) grown for 48h (direct) and from a resuspended plated culture (outgrowth) were immobilized on 0.8% agar pads containing a 1:1000 dilution of PI and imaged as above. If no Δ*mlaE* cells were observed, we replaced the number ‘0’ for quantitative analysis with 0.9, which enables calculation of the CI.

### Flow cytometry analysis of PI-stained cells

Monocultures of WT and Δ*mlaE V. cholerae* were grown for 48h, diluted, and stained with propidium iodide. PI-stained cells were measured on a Sony SH800S cell sorter. Analyses were performed using the FlowJo software (v10.8).

### ATP measurement

ATP measurements were performed using the BacTiter-GloTM Microbial Cell Viability Assay (Promega) following the manufacturer’s recommendations. Briefly, cultures were diluted (1:10) and mixed with an equal volume of the BacTiter-Glo^TM^ Reagent. After 5 min incubation the luminescence was measured using a SpectrMax^R^ i3x plate reader (Molecular Devices).

### OMV purification and quantification

Cultures for OMV quantification were grown at 37°C with shaking at 200 rpm for 8h. After sterile filtration (0.45 µm), the cell free supernatants were incubated with FM4- 64 Dye (5 μg/ml in PBS) (Thermo Fisher Scientific) for 10 min at 37°C (51). After excitation at 515 nm, the emission at 640 nm was measured using a SpectrMax^R^ i3x plate reader. Vesicle production was calculated by dividing relative fluorescence units by OD_600_.

### Transmission electron microscopy

5 µl of WT, Δ*mlaE*, Δ*mlaE fliP*\*, Δ*mlaE fliQ*\* and Δ*mlaE fliF*\* from overnight cultures were adsorbed for 1 minute to a carbon coated grid that had been made hydrophilic by a 20 second exposure to a glow discharge (25mA). Excess liquid was removed with a filter paper; the grid was then floated briefly on a drop of water (to wash away phosphate or salt), blotted again on a filter paper and then stained with 1% Uranyl Acetate for 20-30 seconds. After removing the excess Uranyl Acetate with a filter paper, the grids were examined in a JEOL 1200EX Transmission electron microscope and images were recorded with an AMT 2k CCD camera.

### Motility assay

Motility of *V. cholerae* strains was assessed using swarm agar plates (1% tryptone, 0.5% NaCl, and 0.3% agar). Strains were grown overnight at 37°C on an LB agar plate and a single colony was inoculated by a sterile tip into the swarm plate. The plate was incubated for 20h at room temperature.

### Statistics

Statistical analyses were performed using GraphPad Prism version 10 and are described in each figure legend.

### Software

Data analysis was performed using R and Excel. Graphics and figures were prepared with BioRender, GraphPad Prism and PowerPoint.

## Acknowledgments

We are grateful to members of the Waldor lab for their feedback on this project and manuscript. We are especially grateful to Karthik Hullahalli, Ian Campbell and Yuko Hasegawa for their help with barcode analyses and fruitful discussions. We thank Maria Ericsson at the Harvard Medical School Electron Microscopy Core for assistance with TEM imaging.

This work was supported by NIH grant AI-042347 and the Howard Hughes Medical Institute (HHMI) to M.K.W. and the Life Sciences Research Foundation under grant number: Zingl-2024HHMI to F.G.Z.

This article is subject to HHMI’s Open Access to Publications policy. HHMI lab heads have previously granted a nonexclusive CC BY 4.0 license to the public and a sublicensable license to HHMI in their research articles. Pursuant to those licenses, the author-accepted manuscript of this article can be made freely available under a CC BY 4.0 license immediately upon publication.

## Author contributions

D.R.L. and M.K.W. conceived and all authors designed the study. D.R.L, F.G.Z., A.A.M. and H.Z. and performed all experiments and analyzed data. D.R.L. and M.K.W. wrote the manuscript and all authors edited the paper.

